# A direct high-throughput protein quantification strategy facilitates discovery and characterization of a celastrol-derived BRD4 degrader

**DOI:** 10.1101/2021.12.08.471806

**Authors:** N. Connor Payne, Semer Maksoud, Bakhos A. Tannous, Ralph Mazitschek

## Abstract

We describe a generalizable time-resolved Förster resonance energy transfer (TR-FRET)-based platform to profile the cellular action of heterobifunctional degraders (or proteolysis-targeting chimeras; PROTACs), capable of both accurately quantifying protein levels in whole cell lysates in less than 1 h and measuring small-molecule target engagement to endogenous proteins, here specifically for human bromodomain-containing protein 4 (BRD4). The detection mix consists of a single primary antibody targeting the protein of interest, a luminescent donor-labeled anti-species nanobody, and a fluorescent acceptor ligand. Importantly, our strategy can readily be applied to other targets of interest and will greatly facilitate the cell-based profiling of small molecule inhibitors and PROTACs in high-throughput format with unmodified cell lines. We further-more validate our platform in the characterization of celastrol, a *p*-quinone methide-containing pentacyclic triterpenoid, as a broad cysteine-targeting E3 ubiquitin ligase warhead for potent and efficient targeted protein degradation.

## Introduction

Over the past decade, small molecule-induced selective degradation of specific protein targets has emerged as a novel modality in biomedical research that holds promise to revolutionize drug development, providing access to new classes of therapeutics that can act on disease-relevant proteins previously considered to be undruggable^1, 2^. The concept of targeted protein degradation (TPD) is based on the ligand-induced recruitment of a protein of interest (POI) to an E3 ubiquitin ligase complex for proximity-induced ubiquitination and subsequent proteasomal degradation. Most rational TPD development strategies utilize PROTACs (proteolysis-targeting chimeras), which are heterobifunctional molecules that are formally composed of two individual ligands, one for binding the E3 ligase complex and one for targeting the POI, tethered by a linker^3, 4^.

Despite > 600 E3 ligases in the human proteome that could potentially be exploited, early PROTAC development approaches have focused on a multitude of POIs while relying on only few E3 ligases (e.g. VHL and cereblon) that are targeted by a small number of ligand classes^5, 6^. Although the specific POIs vary between studies, BRD4 (bromo-domain-containing protein 4) is often used as proof-of-concept in PROTAC development programs that explore new E3 ligase binders and/or linker elements^7, 8^. The reason for this is likely a combination of several factors, including historic reference data, abundance, ubiquitous expression, robustness of response, validated orthogonal inhibitor classes, and more recently the availability of commercial building blocks.

Independent of the nature of the POI and the targeted E3 ligase, efficient optimization of PROTACs depends on the availability of robust assay systems that enable the facile, reliable quantification of both biochemical ligand affinities and time- and dose-dependent cellular levels of the POI in response to compound treatment. POI quantification is most commonly done by Western blot analysis, which is inherently time consuming and low throughput. Although various assay technologies, including in-cell Western, enzyme-linked immunosorbent assay (ELISA), AlphaLISA, homogeneous time-resolved fluorescence (HTRF), and luciferase reporter systems, have been developed to increase accuracy and throughput, many depend on the expression of the POI as fusion proteins and/or require expensive specialized equipment and consumables^9^.

To address these shortcomings, we have developed a set of complementary assay strategies based on a common TR-FRET assay platform that greatly facilitates both the characterization of ligand-target engagement, as well as the quantification of endogenous target protein levels directly in cell lysates in high-throughput format. We furthermore employ this approach to identify and characterize celastrol, a triterpene natural product that reversible covalently binds cysteine side chains, as a powerful E3 ligase recruiter for the development of next generation PROTACs.

## Results

### Assay concept and reagent validation

In recent years, assay platforms that combine time-resolved (TR) fluorescence measurements with Förster resonance energy transfer (FRET), often referred to as HTRF immunoassays, have emerged as attractive alternatives to ELISAs and have been successfully employed in PROTAC development for POI quantification and ligand characterization^9–11^. Similar to sandwich ELISAs, HTRF immunoassays generally employ a matched pair of antibodies for POI quantification, yet do not require antibody immobilization or wash steps (Figure S1).

TR-FRET assays are also frequently used to determine the affinity of small molecules for respective POIs (Figure S1). This format generally employs an acceptor-labeled small molecule ligand, referred to as a tracer, in combination with a recombinantly expressed protein featuring an epitope tag (e.g. 6xHis, GST or AviTag) that can be TR-FRET donor-functionalized with a corresponding labeled antibody or streptavidin^12, 13^. While the identification of a linker site for tracer development can be difficult, PROTAC development campaigns by default have solved this problem early on. In fact, several recent independent studies have shown the superior performance of TR-FRET based ligand displacement assays for the characterization of PROTAC binding affinities, kinetics, and ternary complex formation^14–16^.

We hypothesized that rather than utilizing orthogonally labeled matched antibody pairs, which are often difficult to identify, and recombinant POIs, the combination of a tracer with a single antibody directed against the native protein would offer a particularly attractive approach to support PROTAC campaigns by providing a flexible assay platform capable of both ligand affinity profiling and POI quantification directly in cell lysate (Figure 1A). To eliminate the need for direct covalent labeling of the primary antibody, we here developed an approach employing single-domain nanobodies (nano-secondaries), which we labeled with CoraFluor-1, a TR-FRET donor complex with excellent stability and photophysical properties^16^.

**Figure 1.**
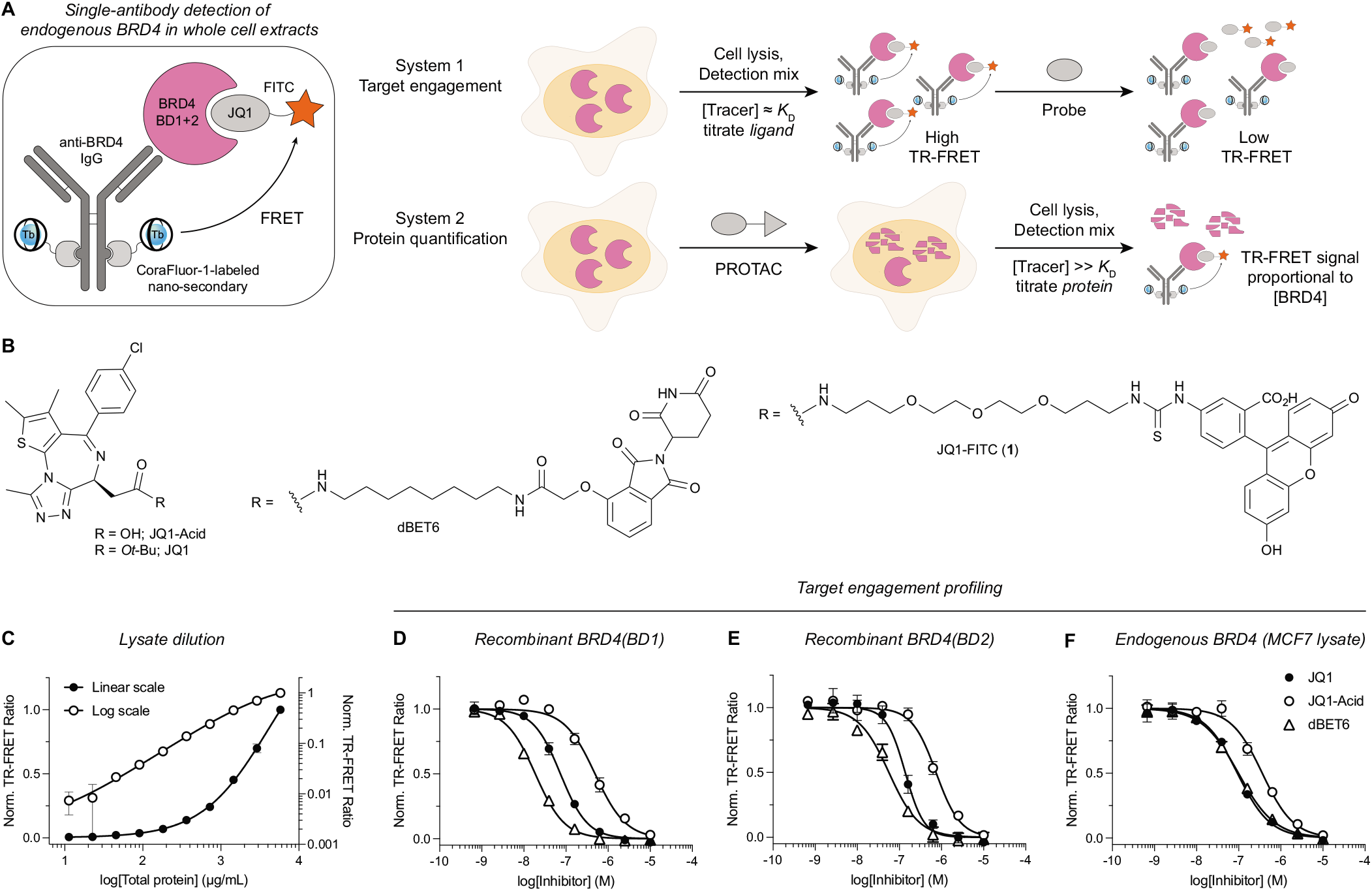
A single-antibody TR-FRET platform to quantitatively measure small molecule target engagement and endogenous protein levels in whole cell extracts. (A) Quantification of both small molecule target engagement and protein levels with endogenous protein targets, here for BRD4. The detection mix consists of a single primary antibody, CoraFluor-1-labeled nano-secondary, and a fluorescent JQ1-based tracer. (B) Chemical structures of bromodomain inhibitors, degraders, and tracers used in this study. (C) TR-FRET-based BRD4 quantification (see Methods) in serially diluted MCF7 cell lysate shows linearity over approximately three orders of magnitude (*n* = 2). (D-F) Dose-titration of small molecule inhibitors and degraders in TR-FRET ligand displacement assays with (D-E) recombinant BRD4(BD1) and BRD4(BD2) domains, and (F) endogenous BRD4 in MCF7 cell extract (0.8 mg/mL total protein) (*n* = 2). Data were fitted to a four-parameter dose-response model in Prism 9. Data in (C-F) are expressed as mean ± SD of *n* technical replicates and are representative of at least two independent experiments.

Because of its extensive use in PROTAC development, we have selected BRD4 for our proof-of-concept studies. Based on the potent prototype BRD4 inhibitor JQ1, we synthesized JQ1-FITC (**1**, Figure 1B) and validated its applicability with individual recombinant bromodomains BRD4(BD1) and BRD4(BD2)^17, 18^. As shown in Figure S2, the tracer potently bound both isolated bromodomains (*K*_D,app_ = 6.5 ± 1.1 nM and 5.8 ± 1.5 nM for BRD4(BD1) and BRD4(BD2), respectively).

### Quantifying BRD4 levels in response to degrader treatment

Following validation of the target engagement assay for recombinant proteins, we next sought to apply the system for the detection of endogenous BRD4. For ligand displacement assays, the tracer is canonically used at or around its *K*_D,app_. In contrast, in protein quantification experiments, the “titration regime” is desired where the tracer concentration is much greater than the *K*_D,app_ to maximize occupancy^19, 20^. In our preliminary experiments, dBET6, a potent BRD4 degrader, was chosen as a positive control due to its well-established activity^21^.

Titration of various lysate dilutions with fixed concentrations of primary anti-BRD4 IgG (0.5 nM), CoraFluor-1-labeled nano-secondary (1 nM), and JQ1-FITC (20 nM) demonstrated linearity over several orders of magnitude with an estimated lower detection limit of ~10 μg/mL total protein (~25 cells/μL) (Figure 1C). Under ligand displacement conditions ([JQ1-FITC] ≈ *K*_D,app_; Figure S2), we determined the *K*_D,app_ values of JQ1, JQ1-Acid and dBET6 toward endogenous BRD4, which closely matched reference data (Figure 1D-F, Table S1)^21^.

Next, under protein titration conditions ([JQ1-FITC] >> *K*_D,app_), we quantified dBET6-induced BRD4 degradation in MCF7 cells. Initially, cells were treated with JQ1 (negative control) or dBET6 (positive control; c_max_ = 10 μM) at varying concentrations for 5 h, followed by a 1 h washout to remove excess ligand^22^. Cells were then lysed in mild lysis buffer (see Methods), followed by the addition of the detection mix (100 nM JQ1-FITC (~11x *K*_D,app_), 0.5 nM anti-BRD4 IgG and 1 nM CoraFluor-1-labeled nano-secondary). Total cell count/protein was achieved by Bradford assay. As expected, we observed a dosedependent decrease in TR-FRET signal in cells treated with dBET6, but not JQ1, indicating potent degradation of BRD4 (DC_50,5h_ = 8.1 ± 1.5 nM; *E*_max,5h_ = 1.1%; Figure 2A and Table S2). The total time from lysis to TR-FRET measurement was ~1.5 h. Western blot analysis on the same samples, which required approximately 2 days, provided near-identical results (Figure 2A).

**Figure 2.**
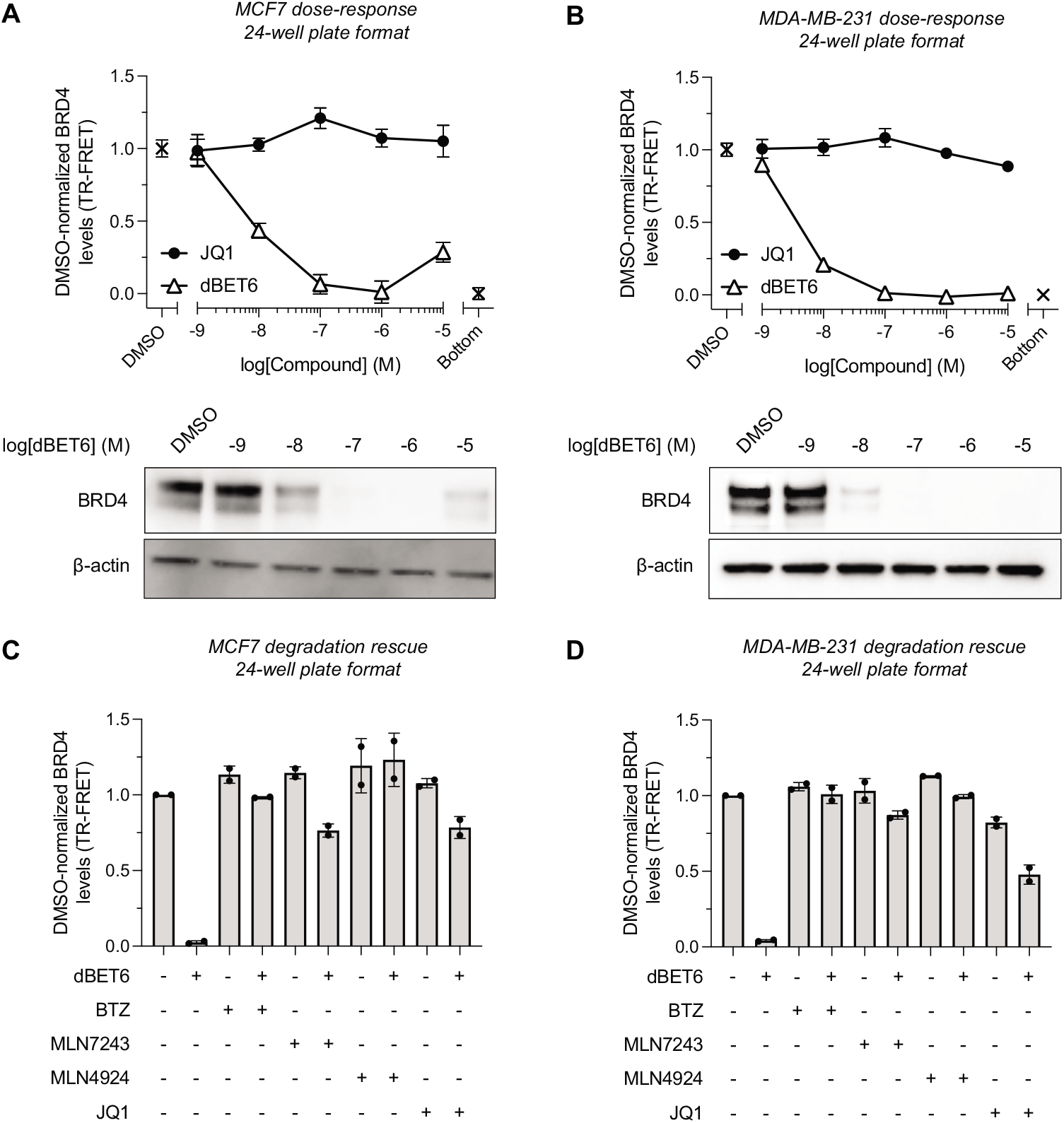
TR-FRET-based quantification of BRD4 levels in unmodified cell lines after degrader treatment. BRD4 protein levels in cell lysate after 5 h treatment with dBET6 (positive control) and JQ1 (negative control) were measured with TR-FRET assay as described in Figure 1A. Assays were run in a 24-well plate format with either (A) MCF7 or (B) MDA-MB-231 cells. Cells were lysed and BRD4 was quantified via addition of TR-FRET detection mix (see Methods). The total time between cell treatment and TR-FRET measurement was |1.5 h. dBET6 showed potent degradation (DC_50,5h,MCF7_ = 8.1 ± 1.5 nM, E_max,5h,MCF7_ = 1.1%; DC_50,5h,MDA-MB-231_ = 4.1 ± 0.3 nM; E_max,5h,MDA-MB-231_ = 1.2%) while JQ1 did not induce BRD4 degradation. Western blot analysis on the same dBET6-treated samples are shown in the bottom panel and are in good agreement with TR-FRET quantification (*n*= 2). (C-D) Quantitative profiling of BRD4 degradation rescue by co-treatment of (C) MCF7 or (D) MDA-MB-231 cells with 1 μM BTZ (20S proteasome), MLN7243 (E1 ubiquitin-activating enzyme), MLN4924 (NEDD8) or 10 μM JQ1 (competing ligand) and 250 nM dBET6 after 5 h shows efficient attenuation of degradation (*n* = 2). Data in (A-D) are expressed as mean ± SD of *n* biological replicates.

Next, to demonstrate compatibility with other cell lines, we performed identical experiments in MDA-MB-231 cells (Figure 2B). Again, dBET6 similarly showed potent BRD4 degradation (DC_50,5h_ = 4.1 ± 0.3 nM; *E*_max,5h_ = 1.2%; Table S2) with good agreement between our CoraFluor TR-FRET platform and Western blot analysis.

We also tested the ability of our TR-FRET assay to quantify the rescue of dBET6-induced BRD4 degradation by bortezomib (BTZ), MLN7243, MLN4924 (1 μM), and JQ1 (10 μM), which constitute 20S proteasome, E1 ubiqui-tin-activating enzyme, NEDD8, and competing inhibitors, respectively, in both MCF7 and MDA-MB-231 cells (250 nM dBET6; Figure 2C-D). In both cell lines, BRD4 degradation was attenuated by all compounds, consistent with previous reports^21^.

### Assay miniaturization to 96-well plate format

PROTAC development and characterization demands the combinatorial analysis of multiple variables including incubation time and compound concentration, which are ideally performed with multiple replicates in parallel to ensure consistency. Accordingly, the number of required data points can quickly grow exponentially. Therefore, rapid, scalable and quantitative assays – especially in unmodified cell lines – are highly desirable^9^. We thus aimed to further miniaturize our assay platform and adapt it to a 96-well plate format, which increases both throughput and compatibility with automated liquid handling equipment. As shown in Figure 3A, upon treatment of MDA-MB-231 cells (20,000 cells/well) with varying concentrations of dBET6 and JQ1 (c_max_ = 1 μM) for 5 h, followed by cell lysis and addition of TR-FRET detection mix (total time to data acquisition = 1 h), we observed a robust dose-dependent decrease in cellular BRD4 levels in dBET6-treated wells (DC_50,5h_ = 3.2 ± 0.1 nM, R^2^ = 0.99; *E*_max,5h_ = 0.6%; Table S2), but not those treated with JQ1.

**Figure 3.**
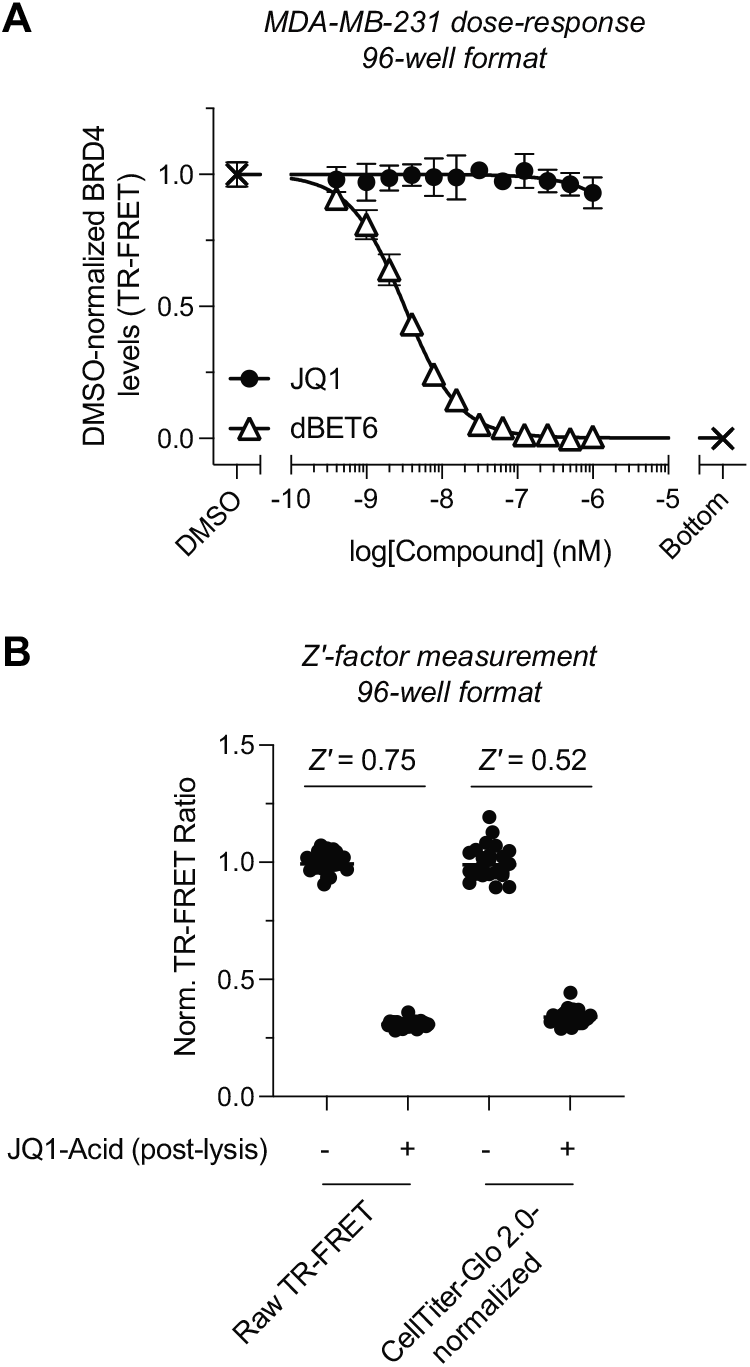
Assay miniaturization and assessment of robustness in 96-well plate format. (A) MDA-MB-231 cells (20,000 cells/well) in 96-well plates were treated with a dose-titration of dBET6 or JQ1 for 5 h. BRD4 levels were quantified via subsequent addition of lysis buffer (60 μL) and detection mix (10 μL) followed by TR-FRET signal acquisition after 1 h incubation (see Methods). dBET6, but not JQ1, induced potent BRD4 degradation (DC_50,5h_ = 3.2 ± 0.1 nM, R^2^ = 0.99, E_max,5h_ = 0.6%). Data are expressed as mean ± SD of *n* = 3 technical replicates. Data were fitted to a four-parameter dose-response model in Prism 9. (B) Z’-factor measurement for TR-FRET quantification assay in 96-well plate format with MDA-MB-231 cells. The Z’-factor was calculated from *n* = 24 positive control wells (DMSO-treated cell lysate) and *n* = 24 negative control wells (DMSO-treated cell lysate + 50 μM JQ1-Acid to simulate 100% BRD4 degradation), with or without CellTiter-Glo 2.0 normalization (also see Figure S3). Data in (A-B) are representative of two independent experiments.

To assess assay performance, we measured the assay robustness which yielded a Z′-factor of 0.75, which is considered excellent for high-throughput screening (HTS) applications (Figure 3B)^23^. Furthermore, to provide an optional mean for data normalization in high-throughput, we employed CellTiter-Glo 2.0 cell viability assay after TR-FRET analysis (Figure 3B)^24^. Since the Z′-factor of CellTiter-Glo 2.0 is 0.83 (Figure S3), normalization results in an overall reduced Z′-factor of 0.52. Regardless, this is still excellent and well suited for HTS.

### Characterization of celastrol-derived BRD4 degrader

Recently, the targeting of other E3 ubiquitin ligase complexes, apart from those canonically used such as CRBN and VHL, using (reversible) covalent ligands that target cysteine side chains has gained increasing attention. Of the predicted > 600 E3 ligases, a substantial fraction feature solvent-exposed cysteine residues that can potentially be exploited with corresponding sulfhydryl-reactive biasing elements, including the recently targeted RNF114, RNF4, FEM1B, DCAF16, and DCAF11 complexes^7, 25^. While selective targeting of E3 ligases can potentially provide cell type- or tissue-specific degradation, we reasoned that targeting multiple ligase complexes with a “promiscuous” reversible covalent inhibitor could provide a means for efficient TPD. The triterpenoid celastrol (CS) can form reversible covalent adducts with multiple cysteine nucleophiles and has been shown to bind a host of proteins^26, 27^. Recently, we have demonstrated that CS also targets Keap1 (Kelch-like ECH-associated protein 1), a redox-regulated member of the CRL3 (Cullin-RING E3 ligase) complex that regulates homeostatic abundance of the transcription factor Nrf2 (Nuclear factor-erythroid factor 2-related factor 2)^16, 28^. CS binds Keap1 BTB and Kelch domains with low micromolar affinity (Table S3). Furthermore, a previous report has demonstrated the promise of PROTACs (CDDO-JQ1) derived from bardoxolone methyl (CDDO-Me), a synthetic triterpenoid that binds Keap1 with high affinity (Figure S4)^29^. Thus, we aimed to evaluate the capacity of CS to function as a recruiting element for E3 ligase activity in a similar manner.

We synthesized a prototype PROTAC candidate CS-JQ1 (**2**, Figure 4A) and validated that CS-conjugation did not impair binding to BRD4 (Figure 4B and Table S1). Interestingly, however, CS-JQ1 lost the ability to bind the Keap1 Kelch domain, while exhibiting slightly improved affinity for the BTB domain (Figure 4C and Table S3). In contrast, we have shown that functionalization of CDDO results in substantially decreased affinity for BTB^16^. Furthermore, we demonstrated that CS-JQ1 was able to induce ternary complex formation between wildtype, fulllength Keap1, and isolated BRD4(BD1) and BRD4(BD2) domains (Figure 4D).

**Figure 4.**
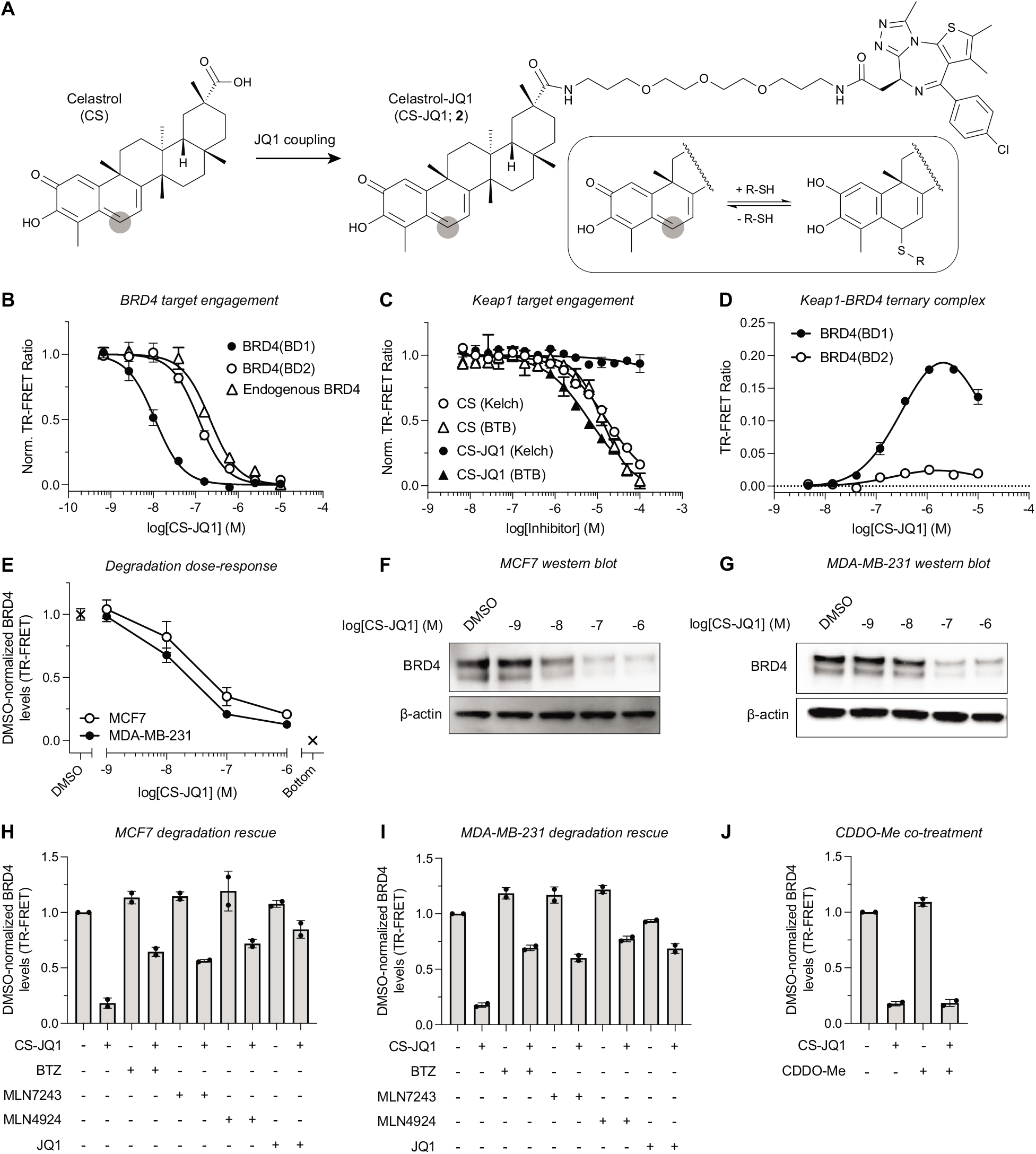
Celastrol is a powerful E3 ubiquitin ligase recruiter for targeted protein degradation applications. (A) Chemical structures of celastrol (CS) and celastrol-JQ1 (CS-JQ1; **2**). Thiophilic sites are highlighted with grey circles. (B) Quantification of target engagement of CS-JQ1 with recombinant BRD4(BD1), BRD4(BD2), and endogenous BRD4 in MCF7 cell extracts (*n* = 2). (C) Dose-titration of CS and CS-JQ1 in TR-FRET assays with full-length Keap1 (see Methods). Data in (B-C) were fitted to a four-parameter dose-response model in Prism 9 (*n* = 2, Kelch; *n* = 4, BTB). (D) CS-JQ1-induced ternary complex formation between full-length Keap1 and BRD4(BD1) and BRD4(BD2) (see methods) (*n* = 2). (E) TR-FRET quantification of BRD4 levels in MCF7 and MDA-MB-231 cells after treatment with dose-titrations of CS-JQ1 for 5 h in 24-well plate assay format. The DC_50,5h_ and E_max,5h_ values for CS-JQ1 were 29 ± 14 (21%) and 16 ± 2 nM (12%) in MCF7 and MDA-MB-231 cells, respectively (*n* =2). (F-G) Western blot analysis of the same samples used for TR-FRET quantification in (E). (H-I) Rescue of CS-JQ1-induced BRD4 degradation (250 nM) by BTZ, MLN7243, MLN4924 (1 μM) and JQ1 (10 μM) in (H) MCF7 or (I) MDA-MB-231 cells (*n* = 2). (J) Co-treatment of MDA-MB-231 cells with CS-JQ1 (250 nM) and potent BTB-targeting Keap1 ligand CDDO-Me (1 μM) does not attenuate BRD4 degradation, indicating potential activity mediated through additional E3 ligase complexes other than Keap1/CRL3 (*n* = 2). Data in (B-D) are expressed as mean ± SD of *n* technical replicates and are representative of at least two independent experiments. Data in (E-J) are expressed as mean ± SD of *n* biological replicates.

Next, MCF7 and MDA-MB-231 cells were treated for 5 h with varying concentrations of CS-JQ1, followed by BRD4 quantification. CS-JQ1 showed dose-dependent and efficient degradation of BRD4 in both cell lines (DC_50,5h,MCF7_ = 29 ± 14 nM; *E*_max,5h,MCF7_ = 21%; DC_50,5h,MDA-MB-231_ = 16 ± 2 nM; *E*_max,5h,MDA-MB-231_ = 12%; Figure 4E and Table S2). Western blot analysis on the same samples yielded results that were again in excellent agreement (Figure 4F-G). We then evaluated 20S proteasomal, E1 ubiquitin-activating enzyme, and NEDD8-dependence on CS-JQ1 activity by co-treatment with 1 μM of either BTZ, MLN7243, MLN4924, respectively (Figure 4H-I). In both cell lines, CS-JQ1-mediated BRD4 degradation (250 nM) was substantially rescued by all three compounds, indicating proteasomal and ubiquitin system dependence. As expected, co-treatment with JQ1 (10 μM) similarly reduced CS-JQ1 degradation efficiency (Figure 4H-I). However, when we evaluated the capacity of CDDO-Me, which we have shown to bind Keap1 BTB domain with potent activity (*K*_D,CDDO-Me_ = 24 nM), to attenuate CS-JQ1-induced BRD4 degradation similar to JQ1, we were surprised to find that CDDO-Me co-treatment did not increase BRD4 levels.

## Discussion

The quantification of cellular protein levels in response to various stimuli represents one of the fundamental experimental techniques employed in chemical biology research. To overcome the inherent limitations of Western blot analysis and ELISA, bioluminescence-based assay formats have been recently developed to simplify workflow, increase throughput, and improve sensitivity. In particular, the use of luciferase tags, such as Promega’s HiBiT system, have gained increasing attention^30^. While these approaches can even enable real-time detection of POI levels in living cells, they are not compatible with unmodified endogenous protein but rather require the expression of the POI as a (split-)luciferase fusion protein. This is usually accomplished by transfection with a corresponding expression vector or by genomic editing of a target cell line. The latter constitutes a lengthy process that generally takes weeks to months and needs to be performed for each cell line of interest.

In contrast, the approach presented here is directly applicable to unmodified cell lines, demonstrates excellent robustness and low technical variability, and is readily adaptable to high-throughput formats, enabling quantitative measurements within 1 h. Unlike luminescence, TR-FRET does not require addition of a luciferase substrate and enzymatic turnover, which provides long signal stability and superior temporal control over assay readout. Another significant improvement is the adaptation of TR-FRET donor-labeled nano-secondaries, which circumvent the need for conjugation of the TR-FRET donor to individual primary antibodies and should find general acceptance for antibody tagging. Furthermore, the monovalent nature of nanobodies avoids the formation of higher order immune complexes common to the use of multivalent secondary detection reagents (e.g. antibodies and streptavidin), which can cause a nonlinear signal response.

The critical prerequisites for our approach are (i) the availability of a primary antibody that recognizes the native POI and (ii) a linker-modified small-molecule ligand that binds the POI with sufficient potency. Both requirements are generally met early on in PROTAC development, which allows for facile implementation of our methodology. Importantly, this approach provides easy access to a ligand displacement platform for the profiling of POI ligands using endogenous protein as a collateral benefit. Owing to the inherent characteristics of TR-FRET, POI-selectivity of the tracer is not required provided that the employed antibody is selective for the POI and does not recognize other proteins to which the tracer also binds.

We further demonstrate the applicability of our TR-FRET quantification approach in the discovery and validation of CS as a promising new ligand for the redirection of E3 ligase activity. Building on our earlier findings and inspired by the previous report of a potent PROTAC derived from the high affinity BTB ligand CDDO and JQ1, we here show that the analogous CS-derived degrader exhibits potent activity in reducing cellular BRD4. Additional mechanistic studies showed proteasomal- and ubiquitin system-dependence, providing support that CS-JQ1 acts through the intended mode of action. Interestingly, co-treatment with CDDO-Me did not reduce the efficacy of CS-JQ1. Since the binding affinity of CS and CS-JQ1 for BTB was determined in a ligand displacement assay using a CDDO-based tracer, it is reasonable to assume that CDDO-Me would efficiently displace CS-JQ1. While we cannot rule out that CS-JQ1 binds Keap1 via a third, previously unrecognized site, it is more likely that CS-JQ1 is dominantly acting by engaging with one or more other E3 ligase complexes. Future studies, which are beyond the scope of this report, are directed towards the identification of the principle E3 ligase(s) exploited by celastrol-based PROTACs.

## Methods

### Mammalian cell culture

MCF7 cells (ATCC) were propagated in RPMI-1640 medium supplemented with 10% FBS, 1% pen-strep, and 1% L-glutamine at 37°C and 5% CO_2_. MDA-MB-231 cells (ATCC) were propagated in DMEM medium supplemented with 10% FBS, and 1%pen-strep at 37°C and 5% CO_2_.

### Preparation of MCF7 cell extracts

A cell pellet from one 15 cm dish (~25 M cells) of MCF7 cells was allowed to thaw on ice and cells were suspended in 400 μL lysis buffer (25 mM HEPES, 150 mM NaCl, 0.2% (v/v) Triton X-100, 0.02% (v/v) TWEEN-20, pH 7.5 supplemented with 2 mM DTT, 250 U Benzonase (Sigma E1014) and 1 mM AEBSF hydrochloride (Combi-Blocks SS-7834)). Optionally, Roche cOmplete, Mini, EDTA-free protease inhibitor cocktail (Sigma 11836170001) can be used in place of, or in combination with, AEBSF hydrochloride. Cells were homogenized via passage through a 27.5-gauge needle 5 times, and the resulting mixture was incubated with slow, end-over-end mixing at 4°C for 30 min. The lysate was clarified via centrifugation at 16,100 x *g* for 20 min at 4°C then 800 μL (1:3 dilution) dilution buffer (25 mM HEPES, 150 mM NaCl, 0.005% (v/v) TWEEN-20, pH 7.5) was added and the lysate was re-clarified at 16,100 × *g* for 20 min at 4°C. Total protein was quantified via detergent-compatible Bradford assay (ThermoFisher 23246) The lysate was used fresh or flash-frozen in liquid nitrogen and stored at −80°C in single-use aliquots.

### TR-FRET measurements

Unless otherwise noted, experiments were performed in white, 384-well microtiter plates (Corning 3572) in 30 μL assay volume. TR-FRET measurements were acquired on a Tecan SPARK plate reader with SPARKCONTROL software version V2.1 (Tecan Group Ltd.), with the following settings: 340/50 nm excitation, 490/10 nm (Tb) and 520/10 nm (FITC) emission, 100 μs delay, 400 μs integration. The 490/10 and 520/10 emission channels were acquired with a 50% mirror and a dichroic 510 mirror, respectively, using independently optimized detector gain settings unless specified otherwise. The TR-FRET ratio was taken as the 520/490 nm intensity ratio on a per-well basis.

### Antibody and nanobody labeling

Nano-secondary alpaca anti-rabbit IgG (ChromoTek shurbGNHS-1), GST V_H_H (ChromoTek st-250), anti-6xHis IgG (Abcam 18184), and anti-GST IgG (Abcam 19256) were labeled with CoraFluor-1-Pfp as previously described^16^. The following extinction coefficients were used to calculate protein concentration and degree-of-labeling (DOL): ChromoTek shurbGNHS-1 *E*_280_ = 24,075 M^−1^cm^−1^, ChromoTek st-250 *E*_280_ = 28,545 M^−1^cm^−1^, IgG *E*_280_ = 210,000 M^−1^cm^−1^, CoraFluor-1-Pfp *E*_340_ = 22,000 M^−1^cm^−1^. Nanobody conjugates were diluted with 50% glycerol and stored at −20°C. IgG conjugates were diluted with 50%glycerol, snap-frozen in liquid nitrogen, and stored at − 80°C.

### Determination of apparent equilibrium dissociation constant (*K*_D,app_) of JQ1-FITC toward individual recombinant bromodomains and endogenous BRD4 in MCF7 lysate

Recombinant BRD4(BD1) and BRD4(BD2) were purchased from BPS Biosciences, Inc and Epicypher, Inc (GST-BRD4(BD1), 31040; GST-BRD4(BD2), 15-0013, respectively). Recombinant bromodomains were diluted to 0.5 nM in assay buffer (25 mM HEPES, 150 mM NaCl, 0.5 mg/mL BSA, 0.005% TWEEN-20, pH 7.5) with 2 nM CoraFluor-1-labeled anti-GST V_H_H, then JQ1-FITC was added in serial dilution (1:1.6 titration, 15-point, c_max_ = 100 nM) using a D300 digital dispenser (Hewlett-Packard) and allowed to equilibrate for 2 h at room temperature before TR-FRET measurements were taken. Nonspecific signal was determined with 50 μM JQ1-Acid, and data were fitted to a One Site – Specific Binding model in Prism 9.

For the profiling of endogenous BRD4, MCF7 cell lysate as prepared above was diluted to 0.8 mg/mL total protein in 1:3 lysis buffer:dilution buffer with 0.5 nM rabbit anti-BRD4 antibody (Cell Signaling Technology; E2A7X) and 1 nM CoraFluor-1-labeled anti-rabbit nano secondary. JQ1-FITC was added in serial dilution (1:1.7 titration, 15-point, c_max_ = 200 nM) using a D300 digital dispenser and allowed to equilibrate for 2 h at room temperature before TR-FRET measurements were taken. Nonspecific signal was determined with 50 μM JQ1-Acid, and data were fitted to a One Site – Specific Binding model in Prism 9.

### TR-FRET ligand displacement assays

The following assay parameters have been used: (i) 4 nM GST-BRD4(BD1), 4 nM CoraFluor-1-labeled anti-GST V_H_H, 20 nM JQ1-FITC in assay buffer, (ii) 4 nM GST-BRD4(BD2), 4 nM CoraFluor-1-labeled anti-GST V_H_H, 20 nM JQ1-FITC in assay buffer, (iii) MCF7 cell lysate at 0.8 mg/mL total protein, 0.5 nM rabbit anti-BRD4 antibody, 1 nM CoraFluor-1-labeled anti-rabbit nano secondary, 20 nM JQ1-FITC. In all cases, test compounds were added in serial dilution (1:4 titration, 9-point, c_max_ = 10 μM) using a D300 digital dispenser and allowed to equilibrate for 2 h at room temperature before TR-FRET measurements were taken. The assay floor (background) was defined with the 10 μM JQ1 dose, and the assay ceiling (top) was defined via a no-inhibitor control.

TR-FRET ligand displacement assays with 6xHis-GST-Keap1 construct were performed as previously described^16^. Data were background corrected, normalized and fitted to a four-parameter dose response model using Prism 9.

### Calculation of inhibitor *K*_D,app_ values from measured TR-FRET IC_50_

For TR-FRET ligand displacement assays with recombinant proteins and for endogenous BRD4 in whole cell extract, we have determined the *K*_D,app_ of the respective fluorescent tracer under each assay condition. Inhibitor *K*_D,app_ values have been calculated using Cheng-Prusoff principles, outlined in Equation 1 below:

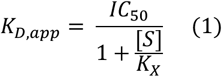

where IC_50_ is the measured IC_50_ value, [S] is the concentration of fluorescent tracer, and *K*_X_ is the *K*_D,app_ of the fluorescent tracer for a given condition^31^.

### TR-FRET assay to measure ternary complex formation between full-length, wildtype Keap1 and individual recombinant bromodomains

Keap1 (tag-free; 11981-HNCB; Sino Biological) was diluted to 40 nM in assay buffer (supplemented with 1 mM DTT) containing 40 nM FITC-Ahx-LDEETGEFL-CONH2 tracer^16^, 20 nM CoraFluor-1-labeled anti-GST antibody, and either 40 nM GST-BRD4(BD1) or GST-BRD4(BD2). CS-JQ1 was added in serial dilution (1:4 titration, 9-point, cmax = 10 μM) using a D300 digital dispenser and allowed to equilibrate for 4 h at room temperature before TR-FRET measurements were taken with identical detector gain settings. Data were background-subtracted from wells containing no CS-JQ1.

### TR-FRET BRD4 quantification assay in 24-well plate format

MCF7 or MDA-MB-231 cells were seeded into 24-well plates (Corning 353047) at a density of 200,000 cells/well in 0.5 mL cell culture medium and allowed to adhere overnight. For dose-response profiling, a D300 digital dispenser was used to dispense serial dilutions of test compounds (1:10 titration, 6-point, c_max_ = 1 or 10 μM) normalized to 0.1% DMSO. Cells were incubated for 5 h at 37°C and 5% CO_2_ then media was replaced with pre-warmed cell culture medium (1 mL/well) and residual test compound was washed out for 1 h at 37°C and 5% CO_2_. After, media was aspirated and cells were washed with PBS (2 mL/well), followed by the addition of ice-cold lysis buffer (200 μL/well). The plate was shaken at 200 rpm on an orbital shaker (IKA KS 260 basic) for 10 min, then lysate was transferred to 1.5 mL Eppendorf tubes and further incubated with slow, end-over-end mixing for 10 min at 4°C. The lysate was clarified via centrifugation at 16,100 × *g* for 10 min at 4°C then total protein concentration was measured using a detergent-compatible Bradford assay (ThermoFisher 23246). Lysate was transferred to a 384-well plate (30 μL × 3 TR-FRET replicates) then 5 μL of 7x detection mix (0.5 nM rabbit anti-BRD4 antibody, 1 nM CoraFluor-1-labeled anti-rabbit nano-secondary, 100 nM JQ1-FITC final concentrations, prepared in dilution buffer) was added to each well and allowed to equilibrate for 1 h before TR-FRET measurements were taken. TR-FRET ratios were background-subtracted from wells containing lysis buffer, 0.5 mg/mL BSA, and detection mix, then normalized to total protein concentration. The average TR-FRET intensity was normalized to DMSO for each biological replicate before being analyzed in Prism 9.

For degradation rescue experiments, a D300 digital dispenser was used to dispense rescue compounds (see respective figure panels for concentrations) normalized to 0.2% DMSO and were pre-incubated for 30 min at 37°C and 5% CO_2_ before degraders (250 nM) were added. Cell treatment, lysate preparation and TR-FRET analysis was performed as described above.

### TR-FRET BRD4 quantification assay in 96-well plate format

MDA-MB-231 cells were seeded into 96-well plates (Corning 3904) at a density of 20,000 cells/well in 100 μL cell culture medium and allowed to adhere overnight. A D300 digital dispenser was used to dispense serial dilutions of test compounds (1:2 titration, 12-point, c_max_ = 1 μM) normalized to 0.1% DMSO. Cells were incubated for 5 h at 37°C and 5% CO_2_ then media was replaced with pre-warmed cell culture medium (150 μL/well) and residual test compound was washed out for 1 h at 37°C and 5% CO2. After, media was aspirated and cells were washed with PBS (200 μL/well), followed by the addition of icecold lysis buffer (60 μL/well). The plate was shaken at 1,000 rpm on an orbital shaker (Boekel Scientific Jitterbug, model 130000) for 10 min, then 10 μL of 7x detection mix (0.5 nM rabbit anti-BRD4 antibody, 1 nM CoraFluor-1-labeled anti-rabbit nano-secondary, 100 nM JQ1-FITC final concentrations, prepared in dilution buffer) was added to each well and allowed to equilibrate for 1 h. The plate was centrifuged at 2,000 × *g* for 1 min then lysate was transferred to a 384-well plate (30 μL × 2 TR-FRET replicates) using an adjustable electronic multichannel pipette (Matrix Equalizer, ThermoFisher 2231) and TR-FRET measurements were taken. Optionally, after TR-FRET measurements, 5 μL/well of CellTiter-Glo 2.0 reagent (Promega G9241) was added to the wells of the 384-well plate and allowed to equilibrate for 10 minutes before luminescence intensity was recorded on a Tecan SPARK plate reader (luminescence module, no attenuation, 250 ms integration time, output Counts/s). TR-FRET ratios were background-subtracted from wells containing lysis buffer, 0.5 mg/mL BSA, and detection mix. The average TR-FRET intensity was normalized to DMSO for each biological replicate, then data were fitted to a four-parameter dose response model using Prism 9.

### Immunoblotting

Proteins in lysates (10 μg) were analyzed by electrophoresis on 3-8% SDS-polyacrylamide gels (ThermoFisher) and subsequently transferred to a nitrocellulose membrane (Bio-Rad). All antibodies were purchased from Cell Signaling Technology. Ponceau staining and β-actin probing (8H10D10, 1:1,000) were used to verify equal protein loading on the blot. The membrane was blocked using 5% nonfat milk powder in TBS-T (Tris-buffered saline; 0.1% TWEEN-20) at room temperature for 1 h and then incubated with an anti-rabbit IgG BRD4 antibody (E2A7X, 1:750) in 2.5% nonfat milk overnight at 4°C. The membrane was then incubated with an anti-rabbit IgG HRP-linked antibody (1:5,000 in 2.5% nonfat milk; 7074S). The proteins were detected using SuperSignal™ West Femto Maximum Sensitivity Substrate (ThermoFisher).

### Reagents and ligands

Reagents and ligands were purchased from Chem-Impex International, Millipore-Sigma, TCI America, Beantown Chemical, Combi-Blocks, MedChemExpress, Ontario Chemicals, and BOC Sciences and used as received.

### Chemical synthesis

Synthetic procedures and small molecule characterization data are provided in Supplementary Note 1.

### Statistics and reproducibility

Biological replicates have been defined as independent cell treatments, performed at different times with biologically distinct samples. For biological replicate analysis, TR-FRET technical replicates refer to the number of replicates performed during the analysis of a given biological sample. For biochemical-based ligand displacement assays, technical replicates refer to the number of parallel replicates used to calculate mean ± SD for a given data point within an experiment. No statistical methods were used to predetermine sample size and investigators were not blinded to outcome assessment.

## ASSOCIATED CONTENT

### Supporting Information

Supplementary Information is linked to this article: Supplementary Tables 1–3, Supplementary Figures 1–4, Supplementary Note 1.

## AUTHOR INFORMATION

### Author Contributions

R.M. conceptualized and supervised all aspects of the project. R.M. and N.C.P. initiated the study. N.C.P. performed all TR-FRET experiments and synthesized compounds. S.M. performed Western blot experiments. R.M., N.C.P., S.M., and B.A.T. designed experiments and analyzed data. R.M. and N.C.P. wrote the manuscript with input from all authors.

## ACKNOWLEDGMENT

This work was supported by NSF 1830941 (R.M.), and NIH 1R01NS064983 (B.A.T. and R.M.). N.C.P. was supported by a National Science Foundation Graduate Research Fellowship (DGE1745303). We thank Dr. Stephen J. Haggarty for generously providing access to tissue culture space. We thank Dr. Jun Qi for donating JQ1-Acid.

## COMPETING INTERESTS STATEMENT

R.M. is a scientific advisory board (SAB) member and equity holder of Regenacy Pharmaceuticals, ERX Pharmaceuticals, and Frequency Therapeutics. R.M. and N.C.P. are inventors on patent applications related to the CoraFluor TR-FRET probes used in this work. R.M., N.C.P., and B.A.T. are inventors on patent applications related to the celastrol-based degraders developed in this work.

## Supplementary Information

**Table S1.** Apparent equilibrium dissociation constants for individual recombinant bromodomains and endogenous BRD4 determined by biochemical TR-FRET ligand displacement assays.

|  | JQ1 |  | JQ1-Acid |  | dBET6 |  | CS-JQ1 |  |
| --- | --- | --- | --- | --- | --- | --- | --- | --- |
| Conditions | $K_{D,app}$ (nM) | 95% CI | $K_{D,app}$ (nM) | 95% CI | $K_{D,app}$ (nM) | 95% CI | $K_{D,app}$ (nM) | 95% CI |
| BRD4(BD1), recombinant<sup>a</sup> | 18 | 17 to 20 | 120 | 100 to 145 | ≤4.6 | 4.1 to 5.1 | ≤2.7 | 2.4 to 3.1 |
| BRD4(BD2), recombinant<sup>b</sup> | 31 | 27 to 36 | 167 | 138 to 202 | 13 | 10 to 16 | 27 | 22 to 32 |
| MCF7 cell lysate, endogenous BRD4<sup>c</sup> | 29 | 25 to 33 | 112 | 93 to 134 | 30 | 28 to 34 | 67 | 56 to 82 |

<sup>a</sup>Conditions: 4 nM recombinant GST-BRD4(BD1), 4 nM CoraFluor-1-labeled anti-GST V<sub>H</sub>H, 20 nM JQ1-FITC, 2 h incubation.
<sup>b</sup>Conditions: 4 nM recombinant GST-BRD4(BD2), 4 nM CoraFluor-1-labeled anti-GST V<sub>H</sub>H, 20 nM JQ1-FITC, 2 h incubation.
<sup>c</sup>Conditions: MCF7 cell lysate at 0.8 mg/mL total protein, 0.5 nM rabbit anti-BRD4 antibody, 1 nM CoraFluor-1-labeled anti-rabbit nano-secondary, 20 nM JQ1-FITC, 2 h incubation.

**Table S2.** Cellular degradation constants for small molecule BRD4 degraders determined by TR-FRET.

|  | dBET6 |  |  | CS-JQ1 |  |  |
| --- | --- | --- | --- | --- | --- | --- |
| Conditions | DC<sub>50,5h</sub> (nM)<sup>a</sup> | 95% CI | $E_{max,5h}$ (%)<sup>b</sup> | DC<sub>50,5h</sub> (nM) | 95% CI | $E_{max,5h}$ (%) |
| MCF7 24-well format | 8.1 | 6.6 to 9.5 | 1.1 | 29 | 15 to 39 | 21 |
| MDA-MB-231 24-well format | 4.1 | 3.8 to 4.5 | 1.2 | 16 | 14 to 18 | 12 |
| MDA-MB-231 96-well format | 3.2 | 3.1 to 3.3 | 0.6 | ND | ND | ND |

<sup>a</sup>DC<sub>50,5h</sub> (half-maximal degradation concentration at 5 h) was determined using a four-parameter dose-response model in Prism 9.
<sup>b</sup> $E_{max,5h}$ (percent remaining target at 5 h) was determined as the lowest remaining percentage of BRD4 signal after background subtraction and DMSO normalization of TR-FRET values.

**Table S3.** Apparent equilibrium dissociation constants for CS and CS-JQ1 toward Keap1-Kelch and Keap1-BTB domains.

| Conditions | CS |  | CS-JQ1 |  |
| --- | --- | --- | --- | --- |
| | $K_{D,app}$ (nM) | 95% CI | $K_{D,app}$ (nM) | 95% CI |
| Kelch<sup>a</sup> | 4,191 | 3,844 to 4,576 | > 100,000 | -- |
| BTB<sup>b</sup> | 1,899 | 1,703 to 2,118 | 946 | 816 to 1,105 |

<sup>a</sup>Conditions: 2 nM 6xHis-GST-Keap1, 0.5 nM CoraFluor-1-labeled anti-6xHis IgG, 5 nM FITC-Ahx-LDEETGEFL-CONH<sub>2</sub> tracer, 4 h incubation.
<sup>b</sup>Conditions: 2 nM 6xHis-GST-Keap1, 0.5 nM CoraFluor-1-labeled anti-6xHis IgG, 40 nM CDDO-FITC tracer, 4 h incubation.

**Figure S1.**
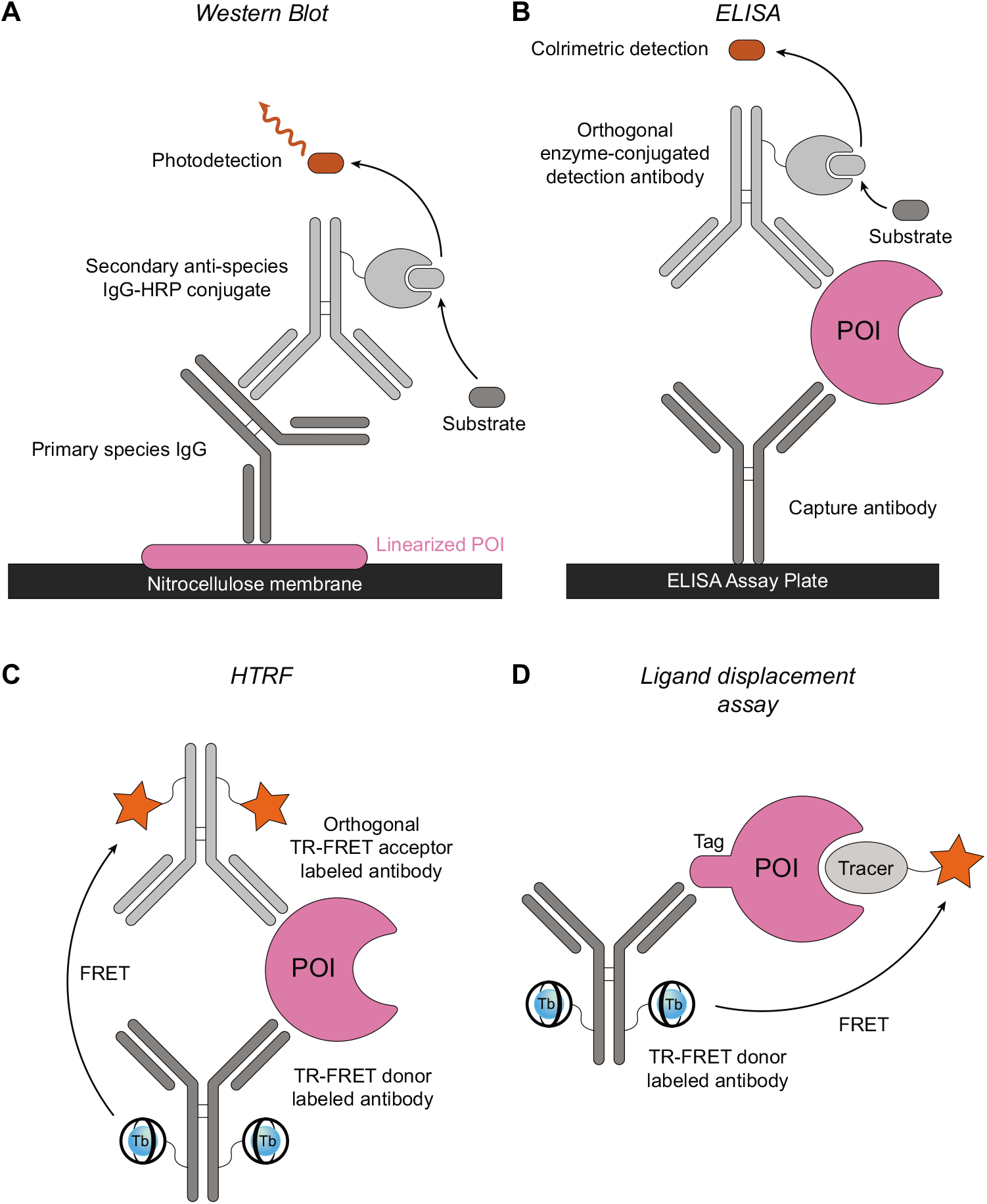
Conventional assay platforms for measuring protein levels and target engagement. (A) Western blot, where proteins are separated by SDS-polyacrylamide gel electrophoresis (SDS-PAGE), transferred to nitrocellulose membranes, and are detected with a primary antibody/HRP-linked secondary system with photodetection as the readout. (B) Sandwich enzyme-linked immunosorbent assay (ELISA). An immobilized capture antibody first binds the POI. After, an enzyme-conjugated detection antibody is added and protein is detected via colorimetric readout. (C) Homogenous time-resolved fluorescence (HTRF) immunoassay. Similar to a sandwich ELISA, orthogonal antibody pairs are used. However, in HTRF antibodies are labeled with a TR-FRET donor and acceptor; concomitant binding to the POI results in an increase in TR-FRET signal. (D) TR-FRET ligand displacement assay. Recombinant, epitope-tagged proteins are incubated with a TR-FRET donor-labeled anti-epitope tag antibody and a fluorescent tracer. Subsequent addition of test compounds displace the fluorescent tracer, resulting in a decrease in TR-FRET signal. Both Western blot and sandwich ELISA-based approaches are generally low-throughput, while ELISA assays are generally more quantitative in nature. In contrast, HTRF-based immunoassays are both quantitative and higher in throughput, yet, like sandwich ELISAs, require matched antibody pairs which are often difficult to obtain.

**Figure S2.**
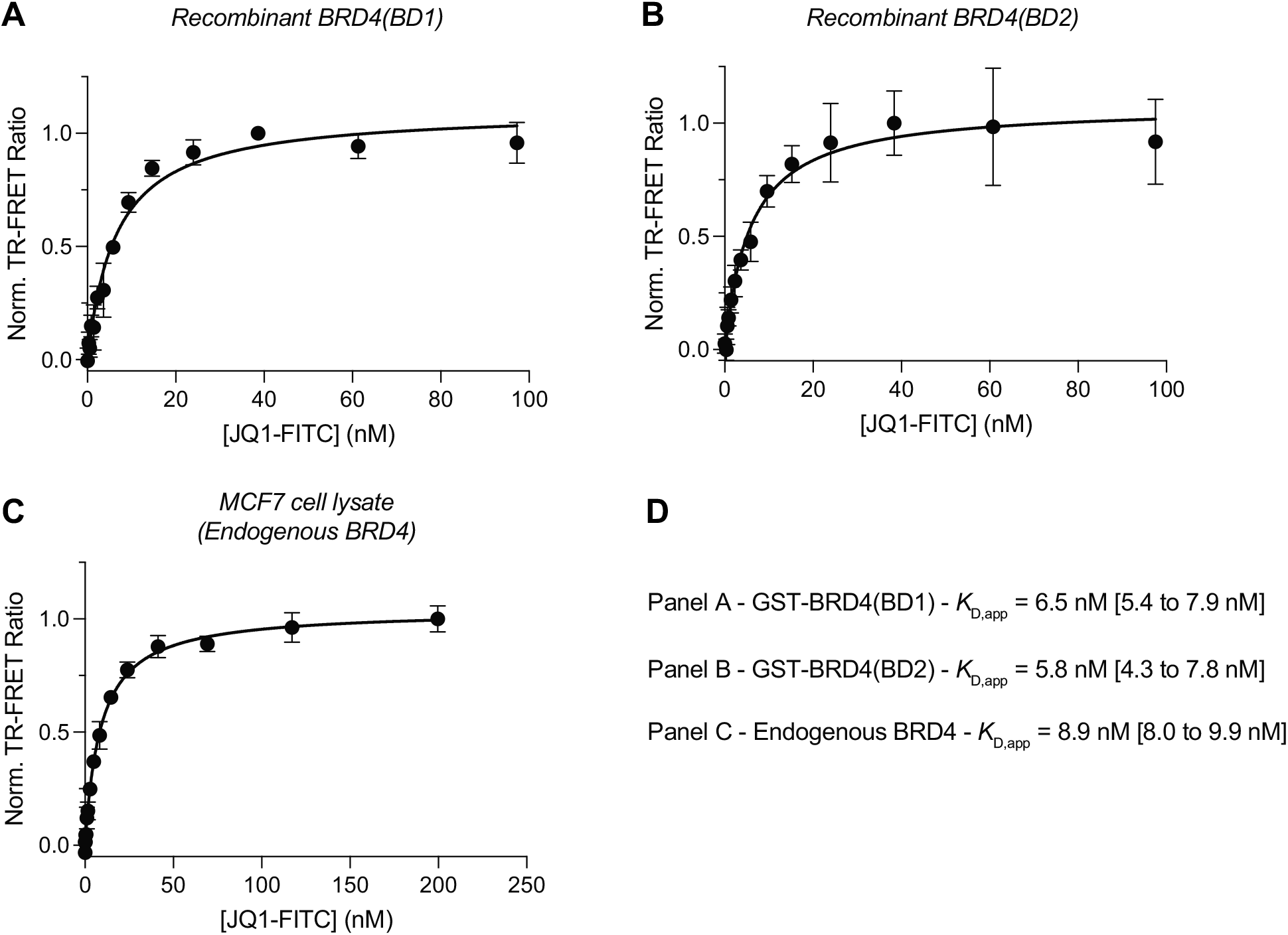
Determination of apparent equilibrium dissociation constant for JQ1-FITC to recombinant bromodomains and endogenous BRD4. Conditions include (A) 0.5 nM GST-BRD4(BD1), 2 nM CoraFluor-1-labeled anti-GST V_H_H, (B) 0.5 nM GST-BRD4(BD2), 2 nM CoraFluor-1-labeled anti-GST V_H_H, (C) 0.8 mg/mL total protein MCF7 lysate, 0.5 nM rabbit anti-BRD4 IgG, 1 nM CoraFluor-1-labeled anti-rabbit nano-secondary (endogenous BRD4). Measured *K*_D,app_ values and associated 95% confidence intervals (shown in parentheses) are displayed in panel (D). Data in (A-C) are expressed as mean ± SD of *n* = 3 technical replicates and are representative of at least two independent experiments. Data were fitted to a one-site model using Prism 9.

**Figure S3.**
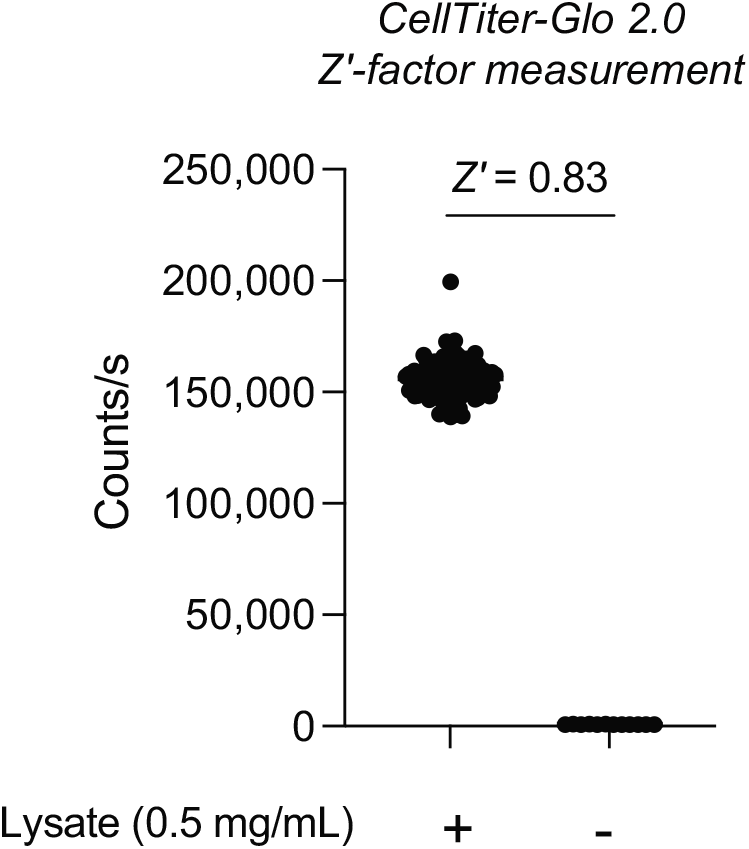
Z′-factor measurement for CellTiter-Glo 2.0. CellTiter-Glo 2.0 reagent (5 μL) was added to either MCF7 cell lysate (0.5 mg/mL; 30 μL; *n* = 96 positive control wells) or lysis buffer containing no cell extract (30 μL; *n* = 12 negative control wells) in a white, 384-well microtiter plate (Corning 3572) and allowed to equilibrate for 10 min at room temperature, after which point luminescence signal was read on a Tecan SPARK plate reader. The Z′-factor is a statistical measure of assay quality using control data, in this case the negative control being lysis buffer in the absence of cell extract (no cellular ATP) and was found to be 0.83, indicating an excellent assay. Data are representative of two independent experiments.

**Figure S4.**
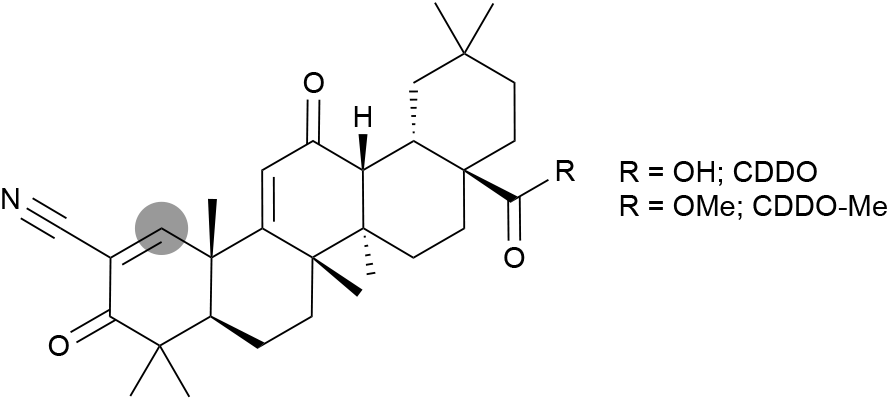
Chemical structures of CDDO and CDDO-Me. Thiophilic site is shown as a grey circle.

## Supplementary Note 1

### Reagents & Equipment

FITC-Ahx-LDEETGEFL-CONH_2_ (FITC-KL9) peptide tracer was custom synthesized by Genscript (Piscataway, New Jersey). CDDO-FITC fluorescent tracer was prepared as previously described (Payne et al., 2021). Column purifications were performed on a Biotage Isolera 4 Purification System equipped with a 200-400 nm diode array detector. For normal phase flash purifications, Sorbtech Purity Flash Cartridges were used (CFC-52300-012-18 and CFC-52500-025-12). For reverse phase flash purifications, Biotage Sfär Bio C18 Duo 300 Å, 20 μm cartridges were used (FSBD-0411-001). Analytical LC/MS was performed on a Waters 2545 HPLC equipped with a 2998 diode array detector, a 2424 evaporative light scattering detector, a 2475 multichannel fluorescence detector, and a Waters 3100 ESI-MS module, using a XTerraMS C18 5 μm, 4.6 x 50 mm column at a flow rate of 5 mL/min with a linear gradient (95% A: 5% B to 100% B 90 sec and 30 sec hold at 100% B, solvent A = water + 0.1% formic acid, solvent B = acetonitrile + 0.1% formic acid). LC/MS data analysis was performed using Waters Masslynx V4.1 SCN 846 software. Proton and carbon nuclear magnetic resonance (^1^H and ^13^C NMR spectra) were recorded on a Bruker Avance III 400 spectrometer using Topspin 3.2 software and data were analyzed using MestreNova (version 12.0.1-20560, Mestrelab Research). Chemical shifts for protons are reported in parts per million (ppm) and are referenced to residual solvent peaks. Data is reported as follows: chemical shift, multiplicity (s = singlet, br s, = broad singlet, d = doublet, t = triplet, q = quartet, p = pentet, m = multiplet), proton coupling constants (*J*, Hz), and integration.

### Synthetic procedures

**5-(3-(1-(4-(4-chlorophenyl)-2,3,9-trimethyl-6*H*-thieno[3,2-f][1,2,4]triazolo[4,3-*a*][1,4]diazepin-6-yl)-2-oxo-7,10,13-trioxa-3-azahexadecan-16-yl)thioureido)-2-(6-hydroxy-3-oxo-3*H*-xanthen-9-yl)benzoic acid – JQ1-FITC – (1)**

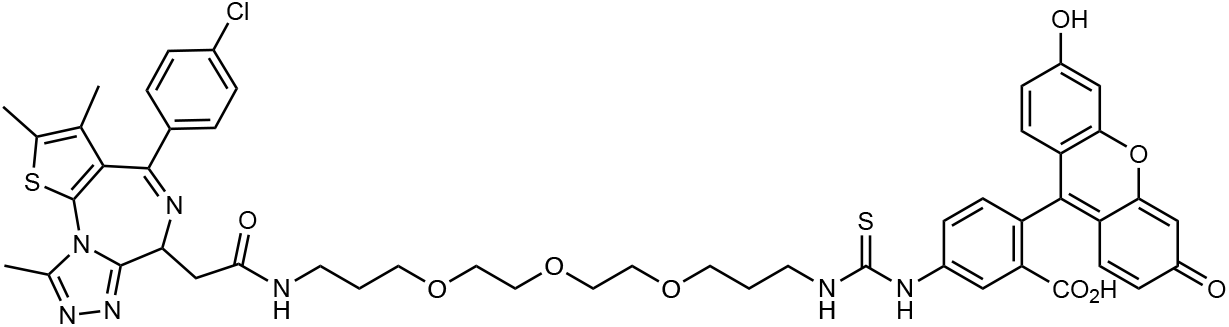

1-(4-(4-chlorophenyl)-2,3,9-trimethyl-6*H*-thieno[3,2-*f*][1,2,4]triazolo[4,3-*a*][1,4]diazepin-6-yl)-2-oxo-7,10,13-trioxa-3-azahexadecan-16-aminium chloride (6.0 mg, 9.4 μmol, 1.1 eq) was dissolved in DMF (200 μL) with DIPEA (15 μL, 11 mg, 85 μmol, 10 eq) then 5(6)-fluorescein isothiocyanate (FITC; 3.3 mg, 8.5 μmol, 1 eq) was added and the reaction mixture was briefly vortexed and allowed to stand at room temperature for 10 min. The reaction mixture was directly purified via reverse phase flash chromatography (λ 220 nm, 250 nm; gradient: 5% ACN/H_2_O + 0.1% formic acid for 3 CV, 5% to 70% ACN/H_2_O + 0.1% formic acid over 14 CV, 70% ACN/H_2_O to 100% ACN + 0.1% formic acid over 1 CV, 100% ACN + 0.1% formic acid for 3 CV). Yield = 4.0 mg, 48% as a yellow solid. ^1^H NMR (400 MHz, DMSO-*d*_6_) δ 10.20 (br s, 3H), 8.42 – 8.32 (m, 1H), 8.30 – 8.15 (m, 2H), 7.75 (d, *J* = 7.7 Hz, 1H), 7.48 (d, *J* = 8.5 Hz, 2H), 7.41 (d, *J* = 8.5 Hz, 2H), 7.16 (d, *J* = 8.3 Hz, 1H), 6.66 (s, 2H), 6.62 – 6.52 (m, 4H), 4.50 (t, *J* = 7.1 Hz, 1H), 3.53 – 3.42 (m, 12H), 3.29 – 3.08 (m, 6H), 2.59 (s, 3H), 2.40 (s, 3H), 1.83 – 1.76 (m, 2H), 1.71 – 1.65 (m, 2H), 1.61 (s, 3H). ^13^C NMR (101 MHz, DMSO-*d*_6_) δ 169.46, 168.58, 163.04, 159.63, 155.12, 151.93, 149.83, 146.83, 141.50, 136.76, 135.24, 132.27, 130.71, 130.14, 129.84, 129.57, 129.07, 128.49, 125.01, 124.06, 112.68, 109.79, 102.24, 69.78, 69.58, 68.20, 68.05, 53.90, 37.65, 35.79, 29.46, 28.58, 14.08, 12.70, 11.32. MS (ESI^+/−^) *m/z* (M+H)^+^ 992.45, *m/z* (M-H)^−^ 990.25, [calculated C_50_H_50_ClN_7_O_9_S_2_: 991.28].

^1^H NMR of compound **1**.

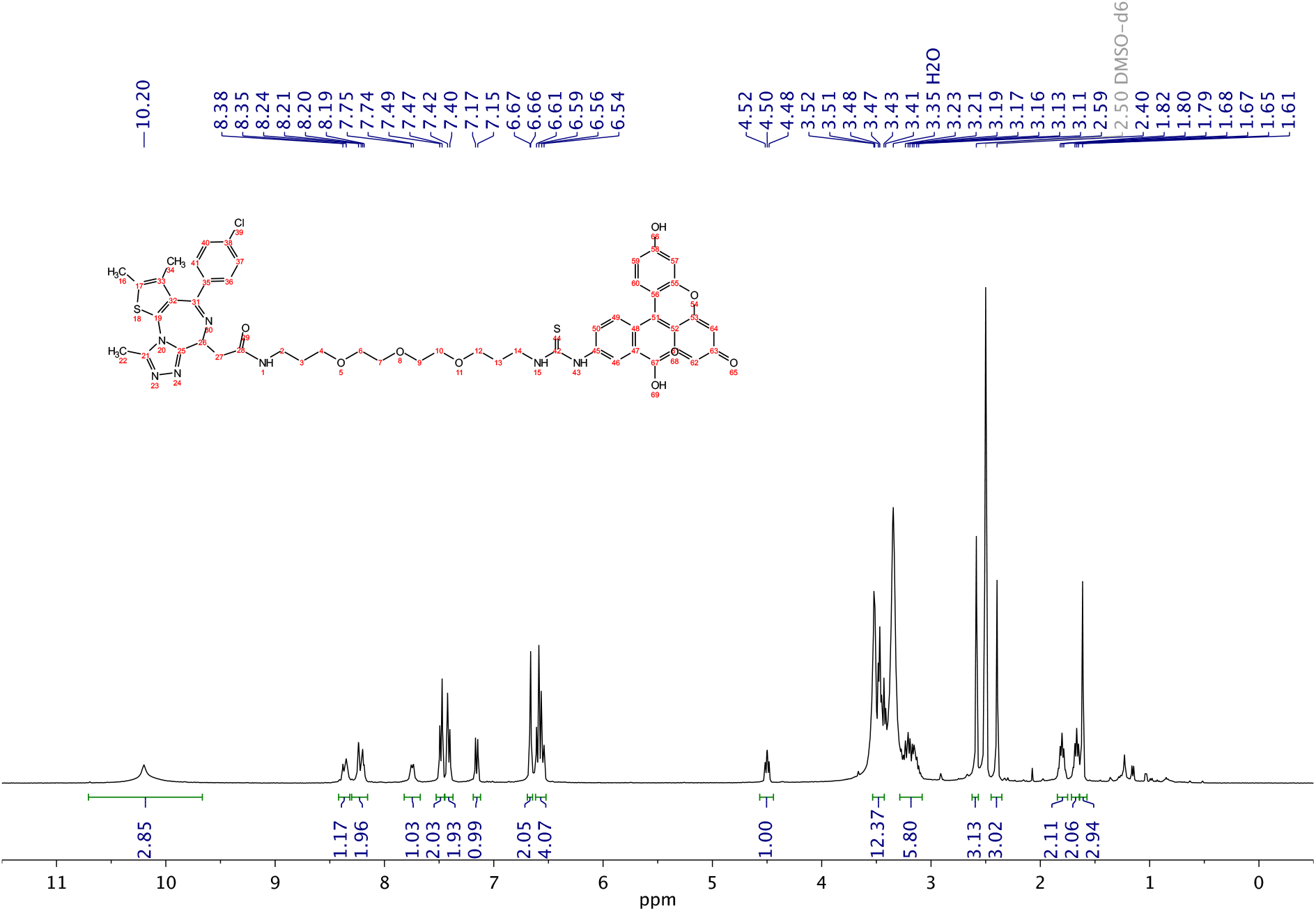

^13^C NMR of compound **1**.

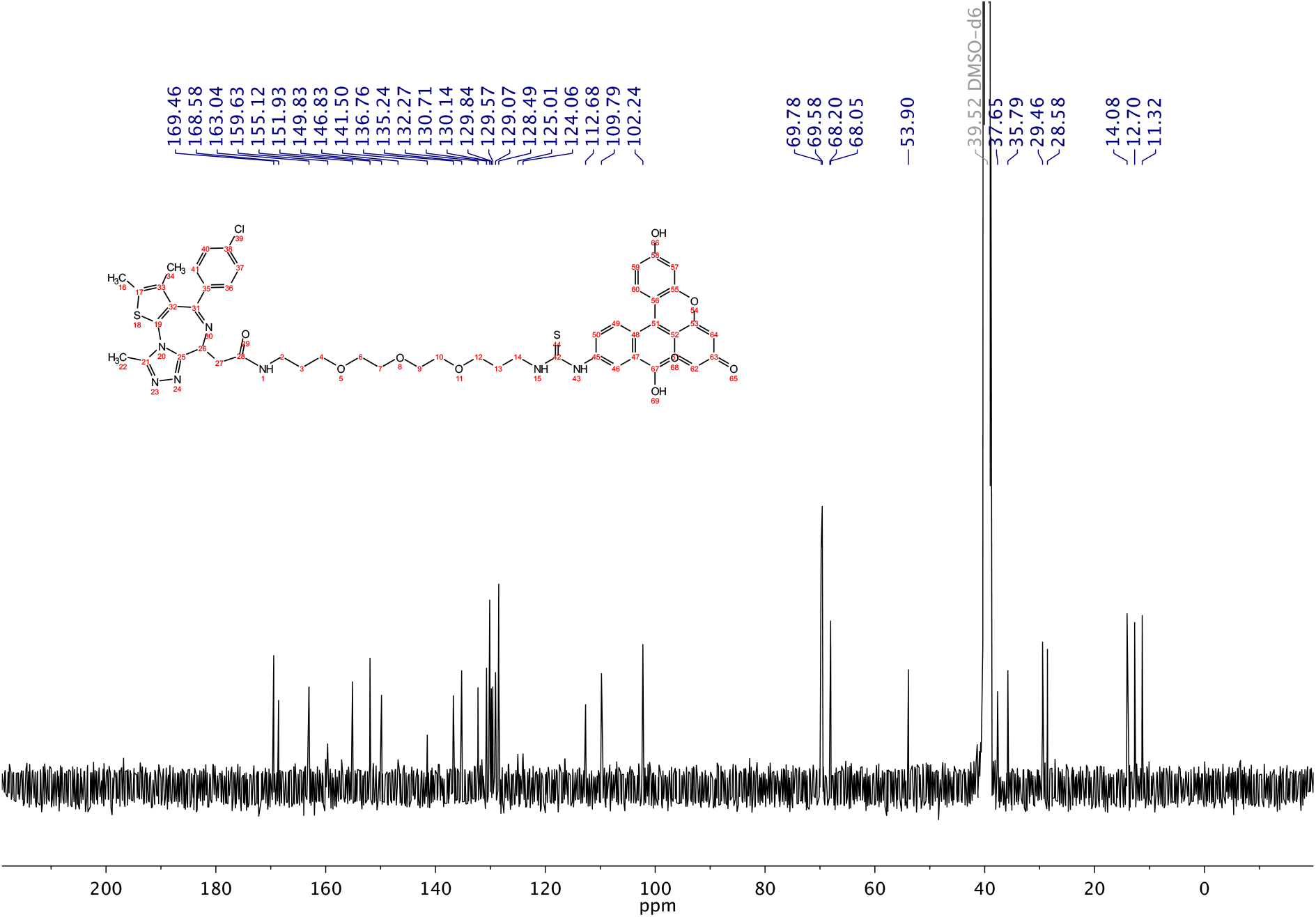

**(2*R*,4a*S*,6a*S*,12b*R*,14a*S*,14b*R*)-*N*-(1-(4-(4-chlorophenyl)-2,3,9-trimethyl-6*H*-thieno[3,2-*f*][1,2,4]triazolo[4,3-*a*][1,4]diazepin-6-yl)-2-oxo-7,10,13-trioxa-3-azahexadecan-16-yl)-10-hydroxy-2,4a,6a,9,12b,14a-hexamethyl-11-oxo-1,2,3,4,4a,5,6,6a,11,12b,13,14,14a,14b-tetradecahydropicene-2-carboxamide – Celastrol-JQ1 – (2)**

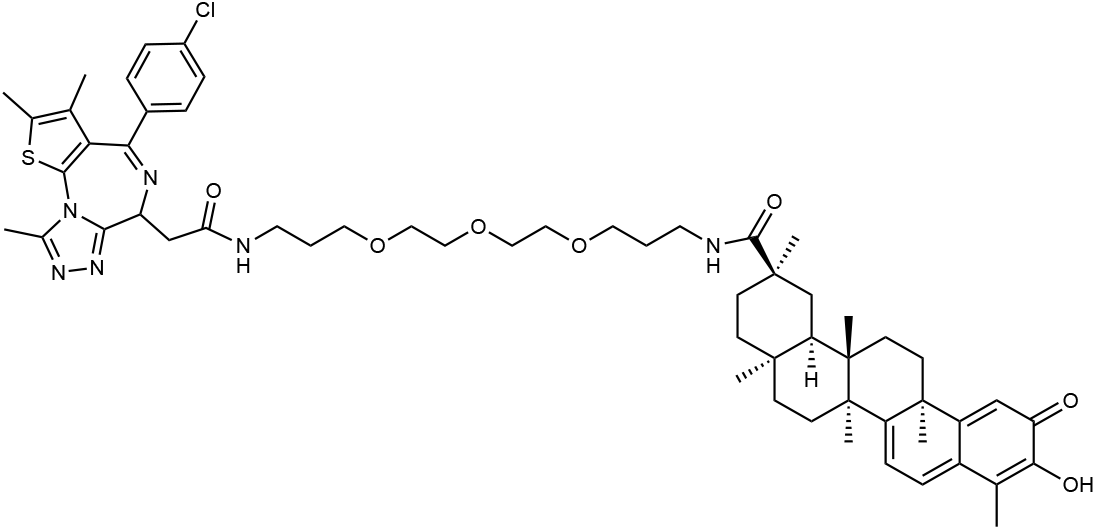

Celastrol (10.0 mg, 22 μmol, 1 eq) was dissolved in DMF (500 μL) then PyBOP (12.1 mg, 23 μmol, 1.05 eq) and 1-(4-(4-chlorophenyl)-2,3,9-trimethyl-6*H*-thieno[3,2-*f*][1,2,4]triazolo[4,3-*a*][1,4]diazepin-6-yl)-2-oxo-7,10,13-trioxa-3-azahexadecan-16-aminium chloride (14.9 mg, 23 μmol, 1.05 eq) were added followed by DIPEA (19 μL, 14 mg, 111 μmol, 5 eq) and the reaction mixture was briefly vortexed then allowed to stand at room temperature for 10 min. The reaction mixture was diluted into EtOAc (50 mL) and the organic layer was washed 2 x equal volume 0.2 N HCl, 2 x H_2_O, 1 x saturated brine solution. The organic layer was then dried over Na2SO4, filtered, and concentrated. The crude product was purified via reverse phase flash chromatography (λ 250 nm, 400 nm; gradient: 5% ACN/H_2_O for 3 CV, 5% ACN/H_2_O to 100% ACN over 18 CV, 100% ACN for 3 CV). Yield = 11.0 mg, 48% as an orange powder. ^1^H NMR (400 MHz, CDCl3) δ 7.40 (d, *J* = 8.2 Hz, 2H), 7.32 (d, *J* = 8.3 Hz, 2H), 7.00 (d, *J* = 6.8 Hz, 2H), 6.88 (t, *J* = 5.0 Hz, 1H), 6.63 (t, *J* = 4.4 Hz, 1H), 6.52 (s, 1H), 6.33 (d, *J* = 7.2 Hz, 1H), 4.64 (t, *J* = 7.0 Hz, 1H), 3.68 – 3.63 (m, 4H), 3.63 – 3.57 (m, 6H), 3.53 (t, *J* = 5.9 Hz, 3H), 3.45 – 3.31 (m, 3H), 3.22 (s, 2H), 2.66 (s, 3H), 2.53 – 2.43 (m, 1H), 2.40 (s, 3H), 2.20 (s, 3H), 2.17 (s, 3H), 2.11 – 1.75 (m, 10H), 1.67 (br s, 6H), 1.59 – 1.48 (m, 5H), 1.42 (s, 3H), 1.12 (s, 3H), 1.09 (s, 3H), 0.63 (s, 3H). ^13^C NMR (101 MHz, CDCl3) δ 178.48, 177.66, 170.61, 170.51, 164.94, 163.95, 155.85, 149.99, 146.14, 136.89, 136.82, 134.24, 132.29, 131.06, 130.92, 130.60, 129.98, 128.84, 127.49, 119.67, 118.12, 117.16, 71.34, 70.65, 70.55, 70.37, 69.96, 54.60, 45.22, 44.55, 43.16, 40.22, 39.53, 39.01, 38.30, 37.97, 36.55, 35.17, 33.92, 33.68, 31.78, 31.23, 31.10, 30.96, 30.12, 29.83, 29.45, 29.18, 28.85, 28.71, 21.88, 18.38, 14.56, 13.25, 12.00, 10.42. MS (ESI^+/−^) *m/z* (M+H)^+^ 1035.79, *m/z* (M-H)^−^ 1033.47, [calculated C_58_H_75_ClN_6_O_7_S: 1034.51].

^1^H NMR of compound **2**.

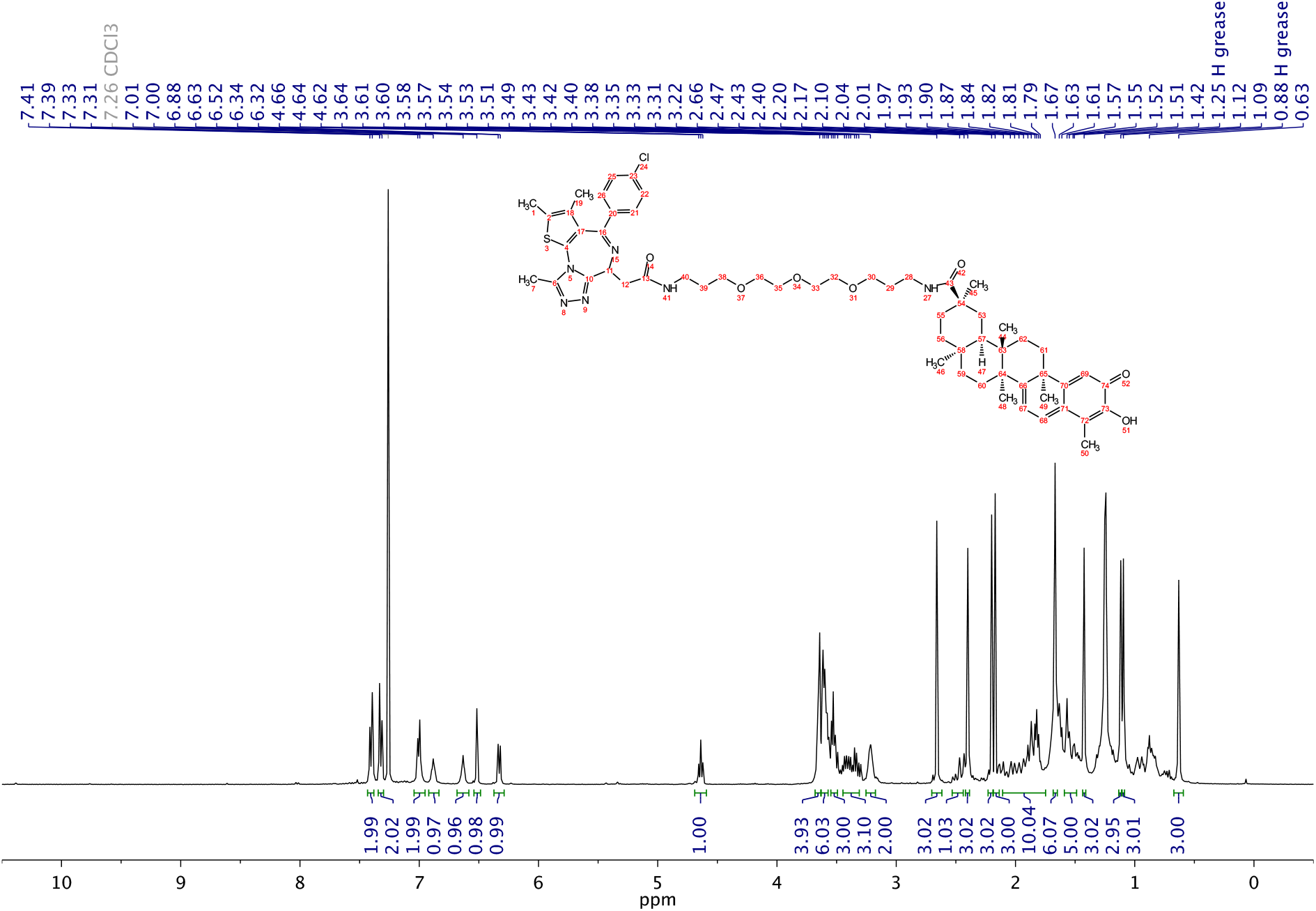

^13^C NMR of compound **2**.

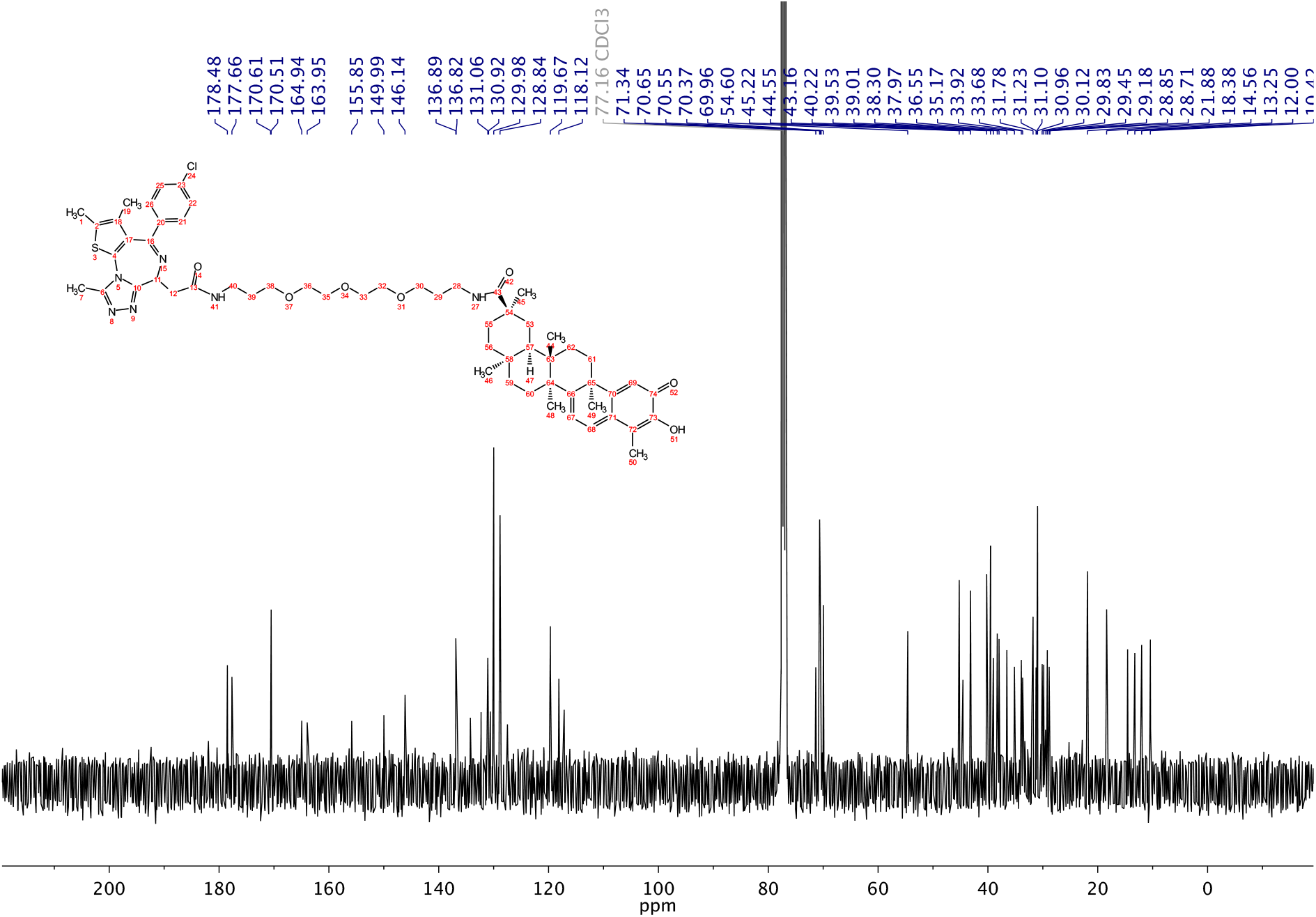

## Notes

### Competing Interest Statement

R.M. is a scientific advisory board (SAB) member and equity holder of Regenacy Pharmaceuticals, ERX Phar-maceuticals, and Frequency Therapeutics. R.M. and N.C.P. are inventors on patent applications related to the CoraFluor TR-FRET probes used in this work. R.M., N.C.P., and B.A.T. are inventors on patent applications related to the celastrol-based degraders developed in this work.

## REFERENCES

1. Burslem, G.M. & Crews, C.M. Proteolysis-Targeting Chimeras as Therapeutics and Tools for Biological Discovery. Cell 181, 102–114 (2020).

2. Chamberlain, P.P. & Hamann, L.G. Development of targeted protein degradation therapeutics. Nat Chem Biol 15, 937–944 (2019).

3. Hanzl, A. & Winter, G.E. Targeted protein degradation: current and future challenges. Curr Opin Chem Biol 56, 35–41 (2020).

4. Maniaci, C. & Ciulli, A. Bifunctional chemical probes inducing protein-protein interactions. Curr Opin Chem Biol 52, 145–156 (2019).

5. Winter, G.E. et al. DRUG DEVELOPMENT. Phthalimide conjugation as a strategy for in vivo target protein degradation. Science 348, 1376–1381 (2015).

6. Buckley, D.L. et al. Targeting the von Hippel-Lindau E3 ubiquitin ligase using small molecules to disrupt the VHL/HIF-1alpha interaction. J Am Chem Soc 134, 4465–4468 (2012).

7. Belcher, B.P., Ward, C.C. & Nomura, D.K. Ligandability of E3 Ligases for Targeted Protein Degradation Applications. Biochemistry (2021).

8. Schapira, M., Calabrese, M.F., Bullock, A.N. & Crews, C.M. Targeted protein degradation: expanding the toolbox. Nat Rev Drug Discov 18, 949–963 (2019).

9. Simard, J.R. et al. High-Throughput Quantitative Assay Technologies for Accelerating the Discovery and Optimization of Targeted Protein Degradation Therapeutics. SLAS Discov 26, 503–517 (2021).

10. Yue, H. et al. Rapid ‘mix and read’ assay for scalable detection of SARS-CoV-2 antibodies in patient plasma. medRxiv (2020).

11. Sy, M., Nonat, A., Hildebrandt, N. & Charbonniere, L.J. Lanthanide-based luminescence biolabelling. Chem Commun (Camb) 52, 5080–5095 (2016).

12. Busby, S.A. et al. Advancements in Assay Technologies and Strategies to Enable Drug Discovery. ACS Chem Biol 15, 2636–2648 (2020).

13. Zwier, J.M. et al. A fluorescent ligand-binding alternative using Tag-lite(R) technology. J Biomol Screen 15, 1248–1259 (2010).

14. Nowak, R.P. et al. Plasticity in binding confers selectivity in ligand-induced protein degradation. Nat Chem Biol 14, 706–714 (2018).

15. Lin, W. & Chen, T. General Stepwise Approach to Optimize a TR-FRET Assay for Characterizing the BRD/PROTAC/CRBN Ternary Complex. ACS Pharmacol Transl Sci 4, 941–952 (2021).

16. Payne, N.C., Kalyakina, A.S., Singh, K., Tye, M.A. & Mazitschek, R. Bright and stable luminescent probes for target engagement profiling in live cells. Nat Chem Biol 17, 1168–1177 (2021).

17. Jung, M. et al. Affinity map of bromodomain protein 4 (BRD4) interactions with the histone H4 tail and the small molecule inhibitor JQ1. J Biol Chem 289, 9304–9319 (2014).

18. Filippakopoulos, P. et al. Selective inhibition of BET bromodomains. Nature 468, 1067–1073 (2010).

19. Jarmoskaite, I., AlSadhan, I., Vaidyanathan, P.P. & Herschlag, D. How to measure and evaluate binding affinities. Elife 9(2020).

20. Wilson, B.D. & Soh, H.T. Re-Evaluating the Conventional Wisdom about Binding Assays. Trends Biochem Sci 45, 639–649 (2020).

21. Winter, G.E. et al. BET Bromodomain Proteins Function as Master Transcription Elongation Factors Independent of CDK9 Recruitment. Mol Cell 67, 5–18 e19 (2017).

22. Donovan, K.A. et al. Mapping the Degradable Kinome Provides a Resource for Expedited Degrader Development. Cell 183, 1714–1731 e1710 (2020).

23. Zhang, J.H., Chung, T.D. & Oldenburg, K.R. A Simple Statistical Parameter for Use in Evaluation and Validation of High Throughput Screening Assays. J Biomol Screen 4, 67–73 (1999).

24. Kepp, O., Galluzzi, L., Lipinski, M., Yuan, J. & Kroemer, G. Cell death assays for drug discovery. Nat Rev Drug Discov 10, 221–237 (2011).

25. Kannt, A. & Dikic, I. Expanding the arsenal of E3 ubiquitin ligases for proximity-induced protein degradation. Cell Chem Biol 28, 1014–1031 (2021).

26. Liu, J., Lee, J., Salazar Hernandez, M.A., Mazitschek, R. & Ozcan, U. Treatment of obesity with celastrol. Cell 161, 999–1011 (2015).

27. Salminen, A., Lehtonen, M., Paimela, T. & Kaarniranta, K. Celastrol: Molecular targets of Thunder God Vine. Biochem Biophys Res Commun 394, 439–442 (2010).

28. Cuadrado, A. et al. Therapeutic targeting of the NRF2 and KEAP1 partnership in chronic diseases. Nat Rev Drug Discov 18, 295–317 (2019).

29. Tong, B. et al. Bardoxolone conjugation enables targeted protein degradation of BRD4. Sci Rep 10, 15543 (2020).

30. Schwinn, M.K. et al. CRISPR-Mediated Tagging of Endogenous Proteins with a Luminescent Peptide. ACS Chem Biol 13, 467–474 (2018).

31. Cheng, Y. & Prusoff, W.H. Relationship between the inhibition constant (K1) and the concentration of inhibitor which causes 50 per cent inhibition (I50) of an enzymatic reaction. Biochem Pharmacol 22, 3099–3108 (1973).

## References

Payne, N.C., Kalyakina, A.S., Singh, K., Tye, M.A., and Mazitschek, R. (2021). Bright and stable luminescent probes for target engagement profiling in live cells. Nature Chemical Biology.

